# Atypical T cell receptor signalling permits distinct gut γδ T cell responses to healthy versus dysregulated tissues

**DOI:** 10.64898/2026.10.09.757810

**Authors:** Leticia Monin, Florencia Cano, James Axford, Grace Cooper, Gerard Hernández-Mir, Klaus Okkenhaug, Adrian C Hayday

## Abstract

Intraepithelial lymphocytes (IEL) are conserved across vertebrates and commonly enriched in γδ T cells. Several such γδ IELs depend upon homeostatic interactions of their T cell receptors (TCRs) with butyrophilin-like (BTNL) proteins expressed by healthy enterocytes and colonocytes. What limits such interactions from inducing full-blown T cell activation which would disrupt barrier integrity is unknown. Here, using primary IEL, we describe an atypical TCR signalling pathway (“signal 1”) downstream of BTNL engagement, that fails to activate either global protein synthesis or cell proliferation. This attenuation could be overcome by provision of a co-stimulator (“signal 2”) or inflammatory cytokines (“signal 3”). It could also be relieved cell-intrinsically by blocking a lipid phosphatase. These findings can, at least in part, explain how extrinsic and intrinsic regulators permit IEL to be constitutively active in healthy tissues and additionally make full-blown responses to adaptive TCR ligands during tissue perturbation.

## Introduction

IEL compose an abundant immunological compartment conserved across gnathostomes and agnathans. However, because of challenges in selectively depleting them vis-à-vis other lymphocytes, their functions are inferred to a greater extent than they have been demonstrated. Nevertheless, the strong consensus is that they contribute to the structural and functional integrity of barrier surfaces. Where studied in appropriate detail, they have displayed at steady-state extremely high levels of cytolytic proteins and chemokines and/or their cognate transcripts (1), with the consequent potential to eradicate neighbouring cells that have become dysregulated and to recruit other immune cells (2,3). Additionally, other effectors may directly regulate epithelial growth and wound-healing (4).

Many IEL compartments are enriched in γδ T cells (5), which are regulated primarily through their T cell receptors (TCRs) (5). In some cases, the TCR repertoire is extremely limited, approaching monoclonal, but in other cases, e.g., mouse and human gut IEL, only one or few TCR-V gene segments may be utilised, but the CDR3 regions are diverse, creating the potential for clonotypic responses (6,7). When viewed from the perspective of conventional T cell biology, this poses a problem, because whereas TCR engagement induces cell proliferation necessary for appropriate clonotypic expansion, unrestrained activation would be structurally disruptive to a barrier surface. Indeed, it is rare for local IEL proliferation to be observed beyond the initial development of the repertoires (2,8,9). This raises the possibility that IEL TCR signal transduction may be distinct from that for conventional T cells.

In this regard, several observations have been reported. First, IEL may be enriched in self-reactive cells that escaped developmental negative selection, perhaps because of attenuated signalling (10). Indeed, it was recently shown that neonatal thymocytes could escape negative selection by virtue of reduced ZAP70 activation downstream of CD3ζ (11) but this has not been examined in IEL.

Second, it has been argued that IEL effector functions are mostly induced by engagement of innate receptors, e.g., NKG2D or IL-12R and IL-18R (12,13). However, it was recently shown that such responsiveness in vivo depends on continued TCR signalling (14). Third, it has been noted that IEL maturation is associated with the acquisition of alternative molecules for TCR signal transduction relative to conventional T cells, e.g., LAT2 (NTAL) in place of LAT (15), UBASH3b instead of UBASH3a (15), SYK instead of ZAP70 (16) and FcεR1γ in place of CD3ζ (14,15). However, the impact of these on IEL TCR signal transduction is unresolved.

Fourth, several studies have characterised gut IEL as exhibiting an “activated yet resting” phenotype because at steady-state they express high levels of effector molecules and/or their transcripts but they do not display fulminant effector function (5). And fifth and finally, it is clear that the TCRs of human and mouse TCRγδ^+^ IEL can engage antigens nonclonotypically via germline-encoded motifs (17–19). Such innate engagement is physiologically important because it mediates TCR-dependent interactions with butyrophilin-like (BTNL) proteins that are expressed by epithelial cells and that are essential for the positive selection and phenotypic maintenance of discrete subsets of TCRγδ^+^ IEL (8,20). Such interactions are not anticipated to induce full-blown effector function that would otherwise perturb the barrier at steady-state. Hence, responses to BTNLs seem highly relevant to the question of how IEL interpret TCR engagement.

To investigate this, we have focussed on primary mouse intestinal IEL. We find that they display strong proliferative responses to agonist anti-CD3 antibodies, showing evidence for largely conventional TCR-dependent signal transduction. Relative to this, TCR engagement of BTNLs transduced atypical and attenuated signals, as did TCR CDR3-dependent engagement of a clonotypic antigen. This attenuation could be relieved by co-stimulation, particularly *via* 4-1BB that was itself upregulated by TCR activation, and by inflammatory cytokines. Given that healthy tissue will not express appreciable levels of co-stimulators or inflammatory cytokines, the homeostatic response to BTNLs will remain attenuated. Additionally, we found that this attenuation could be overridden cell-intrinsically by suppression of a lipid phosphatase and enforced activation of mTOR, thereby identifying a crucial node limiting IEL activation.

## Results

### Qualitatively distinct TCR-dependent responses to BTNL proteins

To assess TCR-mediated responses of intestinal IEL, we used primary cells from mouse small intestines. Among these are γδ T cells expressing TCR Vγ7, which mediates responses to BTNL1+BTNL6 expressed by epithelial cells. Despite their commonly reported refractoriness to stimulation, primary IEL responded strongly to anti-CD3 antibodies, by comparison to control immunoglobulin (Ig), as measured by significant upregulation of activation markers, including CD25 (the IL2Rα chain), CD71 (the transferrin receptor) and 4-1BB (CD137)(Figure 1a). Albeit to a lesser extent, this was also true for Vγ7^+^ IEL responding to murine intestinal epithelial MODE-K cells transduced with a vector expressing BTNL1+BTNL6 but not to MODE-K cells transduced with an empty vector (EV; Figure 1a). This was accompanied by increased eGFP expression in NR4A1 (*Nur77*)-eGFP fluorescent reporter mice(21), indicating direct TCR signalling (Figure 1b). Of note, BTNL1+6 activation was specific for Vγ7^+^ cells and did not elicit TCRβ^+^ IEL activation in the same cultures (Figure 1b). Moreover, the eGFP signal was comparable for anti-CD3 and BTNL1+6, possibly suggesting that the lower activation marker responses might reflect selective attenuation downstream of BTNL engagement by the TCR, an issue that is further explored below.

**Figure 1.**
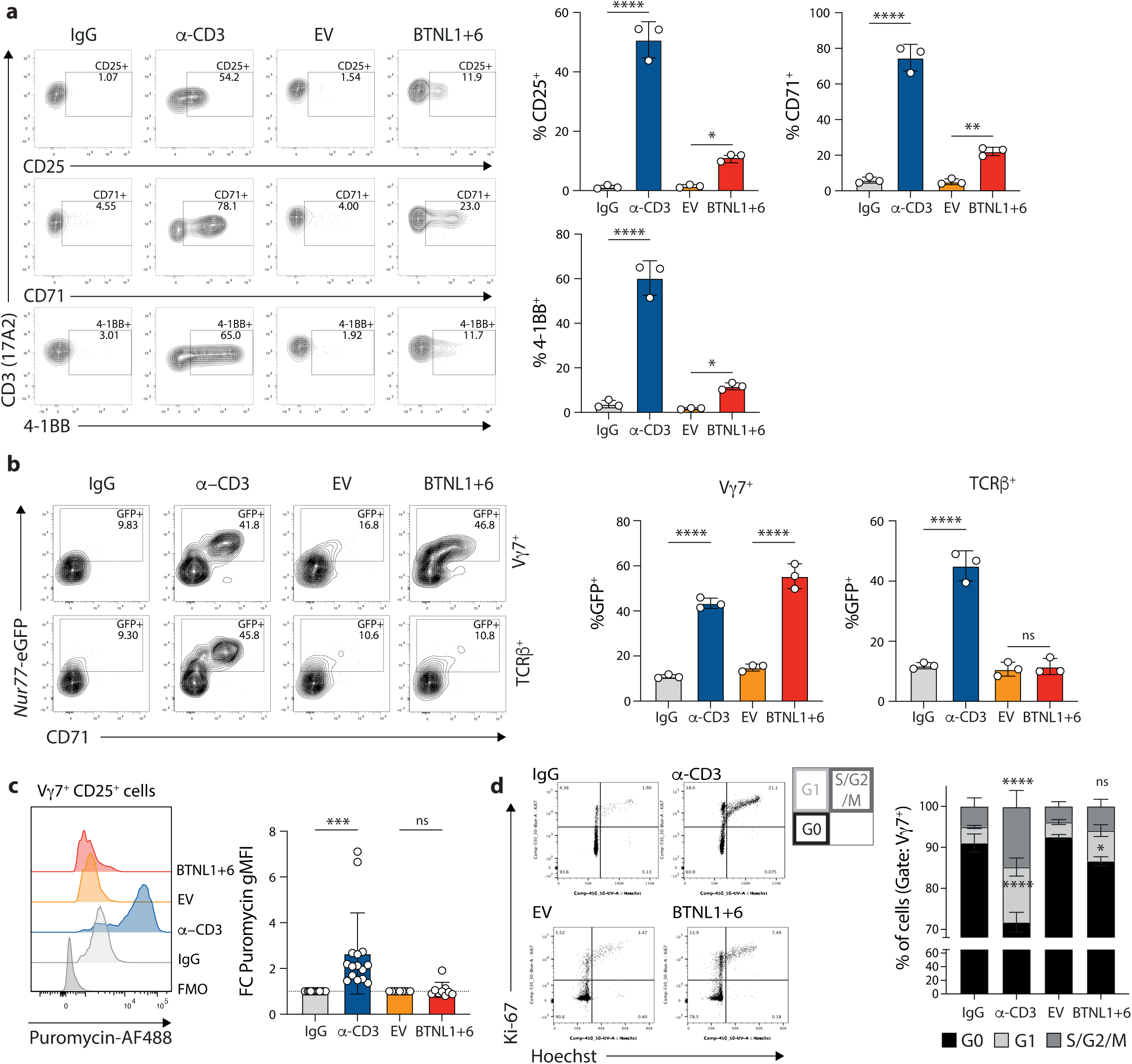
**a.** WT IEL were co-cultured o.n. with plate-bound IgG or anti-CD3, or with MODE-K cells transduced with empty vector (EV) or BTNL1+6. IEL activation was measured by analysing CD25, CD71 and 4-1BB on Vγ7+ cells. **b.** Nur77 expression downstream of IgG/anti-CD3 or MODE-K EV/BTNL1+6 was evaluated in Vγ7+ or TCRβ+ cells by flow cytometry after o.n. stimulation. **c.** Global translation was analysed in activated, stimulated IEL by puromycing labelling using flow cytometry. **d.** Analysis of cell cycle in activated Vγ7+ cells using Ki-67 and Hoechst staining.

Following anti-CD3 engagement, gut Vγ7^+^ IEL increased both CD25 expression (Supp Fig 1) and overall protein translation rates, as measured by ex vivo puromycin incorporation (Figure 1c). This was accompanied by another typical T cell response to TCR engagement, namely increased cell cycle entry: whereas less than 10% of primary Vγ7^+^ IEL were cycling, anti-CD3 stimulation saw ∼30% of cells enter G1 or S-G2/M (Figure 1d). In stark contrast, activated BTNL-stimulated Vγ7^+^ IEL showed no aggregate increase in protein translation (Figure 1c). Whereas there was a small but significant increase in Vγ7^+^ IEL progressing into G1 as well as CD25 upregulation in matched samples (Supp Fig 1), there was no significant increase in cells entering S-G2/M (Figure 1d). These data highlight that although both stimuli engage the TCR (comparable eGFP upregulation), BTNL1+6 stimulation fails to trigger full anabolic and proliferative reprogramming in gut IELs.

We next sought to identify any divergence in TCR signal transduction that might be associated with the difference in response to anti-CD3 and BTNL1+6. Given the challenges of handling primary IEL, phospho-flow analyses of signal transduction gave predictably variable results, but across a number of studies the data clearly showed that both anti-CD3 and BTNL1+6 triggered activation of proximal TCR signalling components, including CD3ζ phosphorylation, reduced phosphorylation of both the inhibitory residue in LCK, degradation of the inhibitory IκB molecule, and downstream phosphorylation of ERK (Figure 2a). Moreover, the specificity of BTNL signalling for Vγ7^+^ IEL was evident by its negligible impact on TCRαβ^+^ IEL (Figure 2a). However, in contrast with anti-CD3, BTNL1+6 failed to activate key signalling and metabolic regulators, including ZAP70, SYK, AKT and mTOR (Figure 2a).

**Figure 2.**
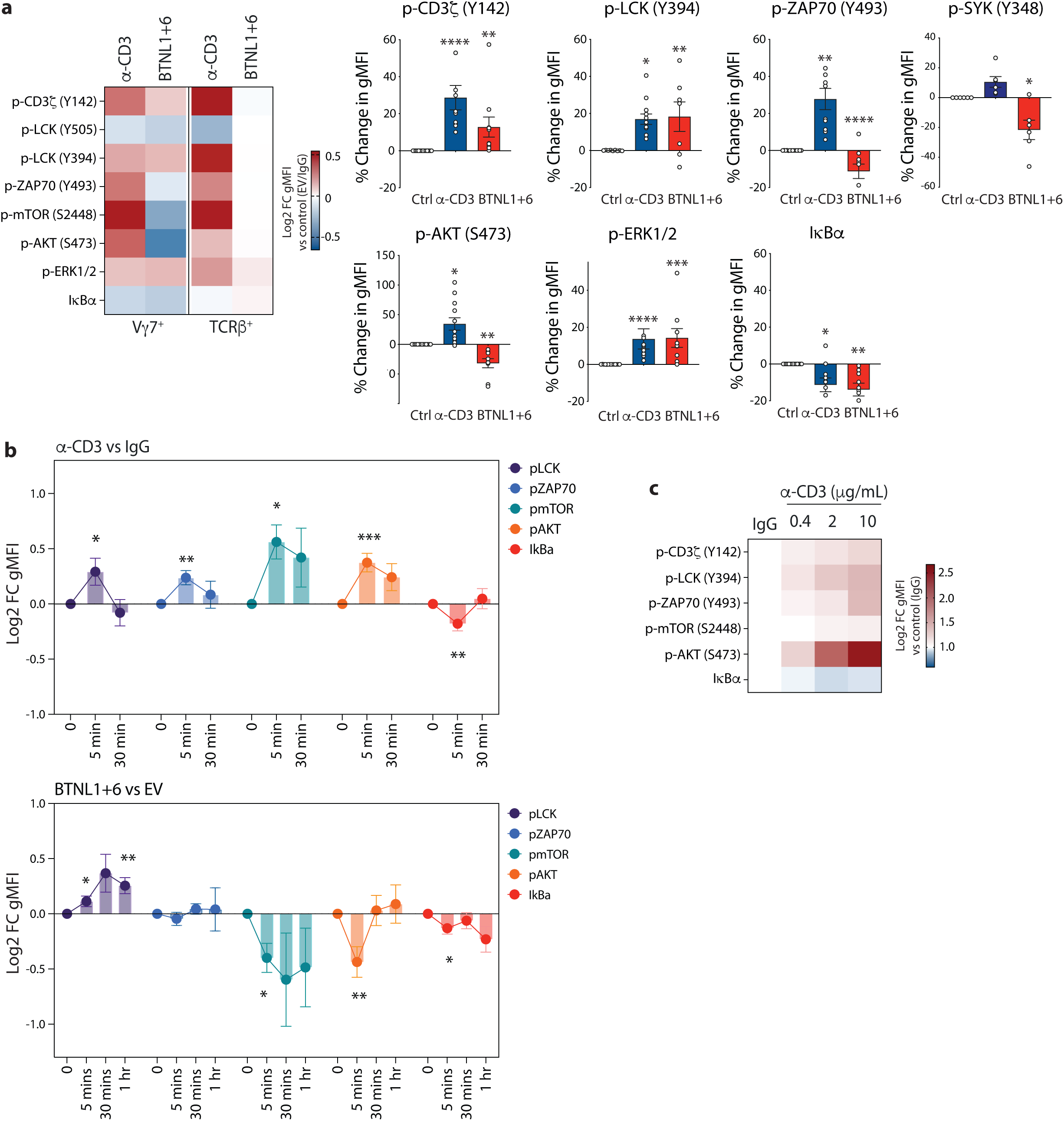
**a.** WT IEL were stimulated with plate-bound IgG or anti-CD3, or with MODE-K cells transduced with empty vector (EV) or BTNL1+6. Signalling pathway activation was analysed by phospho-flow 5 minutes post-stimulation. **b.** Time-course of p-LCK, p-ZAP70, p-mTOR, p-AKT and IkBa degradation following stimulation. **c.** Dose-response of signalling pathway activation following IgG and anti-CD3 stimulation. In all cases, values show the fold change of activation relative to the IgG control (for anti-CD3) or MODE-K EV (for BTNL1+6).

Although AKT and mTOR activation are typically sustained by phosphoinositide3-kinase (PI3K) activated downstream of T cell co-stimulation (a.k.a. Signal 2) (22), TCR signal alone (a.k.a. Signal 1) is typically sufficient to cause transient AKT and mTOR activation. We therefore investigated whether the relative failure of BTNL1+6 to activate this pathway reflected different kinetics and/or strengths of the activation signal. However, neither was the case, since a longitudinal analysis of TCR signalling intermediates revealed sustained divergence in mTOR and AKT phosphorylation across all timepoints (Figure 2b), while titration of the anti-CD3 concentration resulted in graded reductions but not reversal of mTOR and AKT activation (Figure 2c). Thus, under the conditions employed, BTNL1+6 and anti-CD3 delivered related but qualitatively distinct TCR signals.

### Mostly similar IEL responses to innate versus adaptive γδTCR ligands

Having established the competence of primary γδ IEL to respond to TCR engagement via either anti-CD3 or BTNL1+6, we wished to compare the TCR-mediated response to BTNL1+6 with the response to a cognate ligand bound via the TCR CDR3 loops. Whereas the former type of engagement typically describes homeostatic IEL interactions with healthy cells, the latter typically describes effector-inducing interactions with so-called “stress antigens” which are over-expressed or mis-expressed on infected or transformed cells (23). One such ligand is the MHC-related T10/T22, binding to which is mediated primarily by CDR3δ sequences (24,25). Those specific CDR3δ sequences are estimated to emerge through gene rearrangement in less than 1% of TCRγδ^+^ IEL (26), which contrasts greatly with the response to BTNL1+6 shown by essentially all Vγ7^+^ IEL (18). Importantly, we previously characterised a Vγ7Vδ7 IEL TCR reactive both to BTNL1+6 and T10/T22(18), and we therefore used this TCR to establish a congenically marked retrogenic mouse system (27) allowing us to obtain large numbers of primary γδ T cells with this defined dual-reactivity (Figure 3a). Immune reconstitution, homing and surface immunophenotype of the developing γδ T cells were assessed ∼6-8 weeks after transplantation.

**Figure 3.**
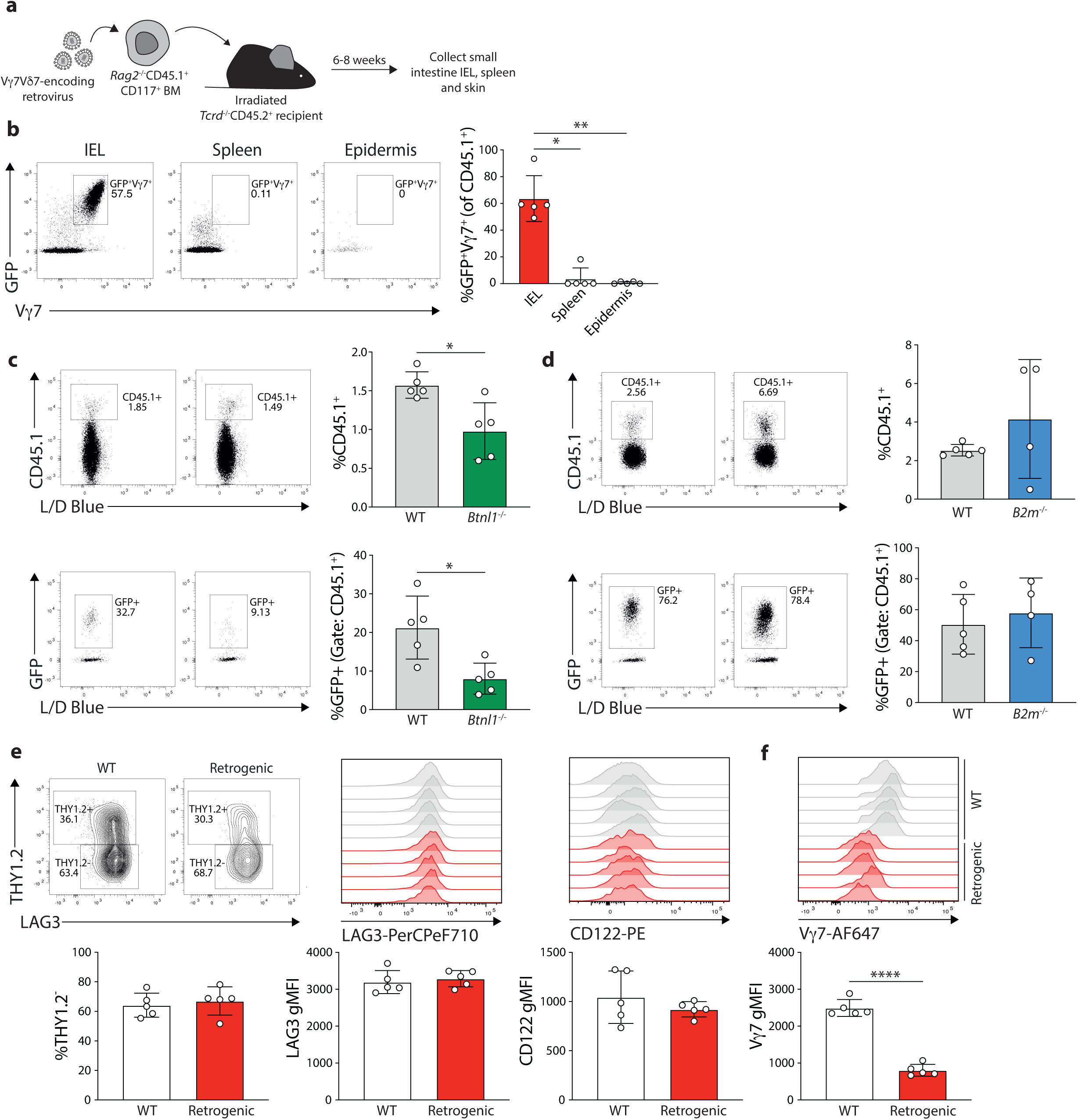
**a.** Experimental setup for retrogenic mouse generation. MACS-sorted CD117+ bone marrow progenitors from *Rag2*^-/-^CD45.1^+^ mice were transduced with a Vγ7Vδ7 TCR-encoding retrovirus. *Tcrd*^-/-^CD45.2^+^ recipients were irradiated and adoptively engrafted with the transduced cells the following day. Mice were humanely killed 6-8 weeks post-transfer and tissues harvested for analysis. **b.** Flow cytometric analysis of Vγ7 and GFP expression in the IEL compartment, spleen and epidermis of retrogenic mice (Gate: CD45.1+ cells). Developmental requirements of retrogenic cells were tested by generating retrogenics in *Btnl1*^-/-^ (**c**) *B2m*^-/-^ (**d**) recipients. The maturation status (**e**) and TCR expression level (**f**) of retrogenic cells was compared to that of their WT counterparts by flow cytometry, using WT CD45.2^+^ mice as recipients.

Retrogenic Vγ7Vδ7 T cells preferentially localised to the intraepithelial intestinal compartment, being, for example, essentially absent from the epidermis and spleen, as well as from the intestinal lamina propria (Figure 3b; Supp Fig 2), results that would be consistent with a developmental dependence on intestinal epithelial-specific interactions with BTNL proteins. Indeed, whereas CD45.1^+^ donor cells could be detected in *B2m*^-/-^and *Btnl1*^-/-^ recipients, there was a significantly reduced capacity to identify GFP-labelled Vγ7Vδ7 IEL in *Btnl1*^-/-^ mice (Figure 3c), whereas IEL reconstitution was unaffected in *B2m*^-/-^ recipients, reflecting the cells’ independence from MHC Class I (Figure 3d).

The intestinal retrogenic cells exhibited a mature IEL immunophenotype (8), as illustrated by over 60% of cells showing low THY1.2 expression and high expression of LAG3 and CD122 (Figure 3e); however, they did also exhibit reduced surface TCR expression (Figure 3f). We next compared retrogenic IEL activation in response to BTNL1+6-expressing *versus* T22-expressing MODE-K cells. γδ IEL from retrogenic mice, but not their αβ counterparts, responded to both stimuli, as evidenced by increased CD25 upregulation (Figure 4a). Despite this activation, neither BTNL1+6 nor T22 increased global protein translation or (Figure 4b) or cell cycle transition to S-G2/M (Figure 4c), in contrast with the robust translational and proliferative responses elicited by anti-CD3 (Figure 4b, c).

**Figure 4.**
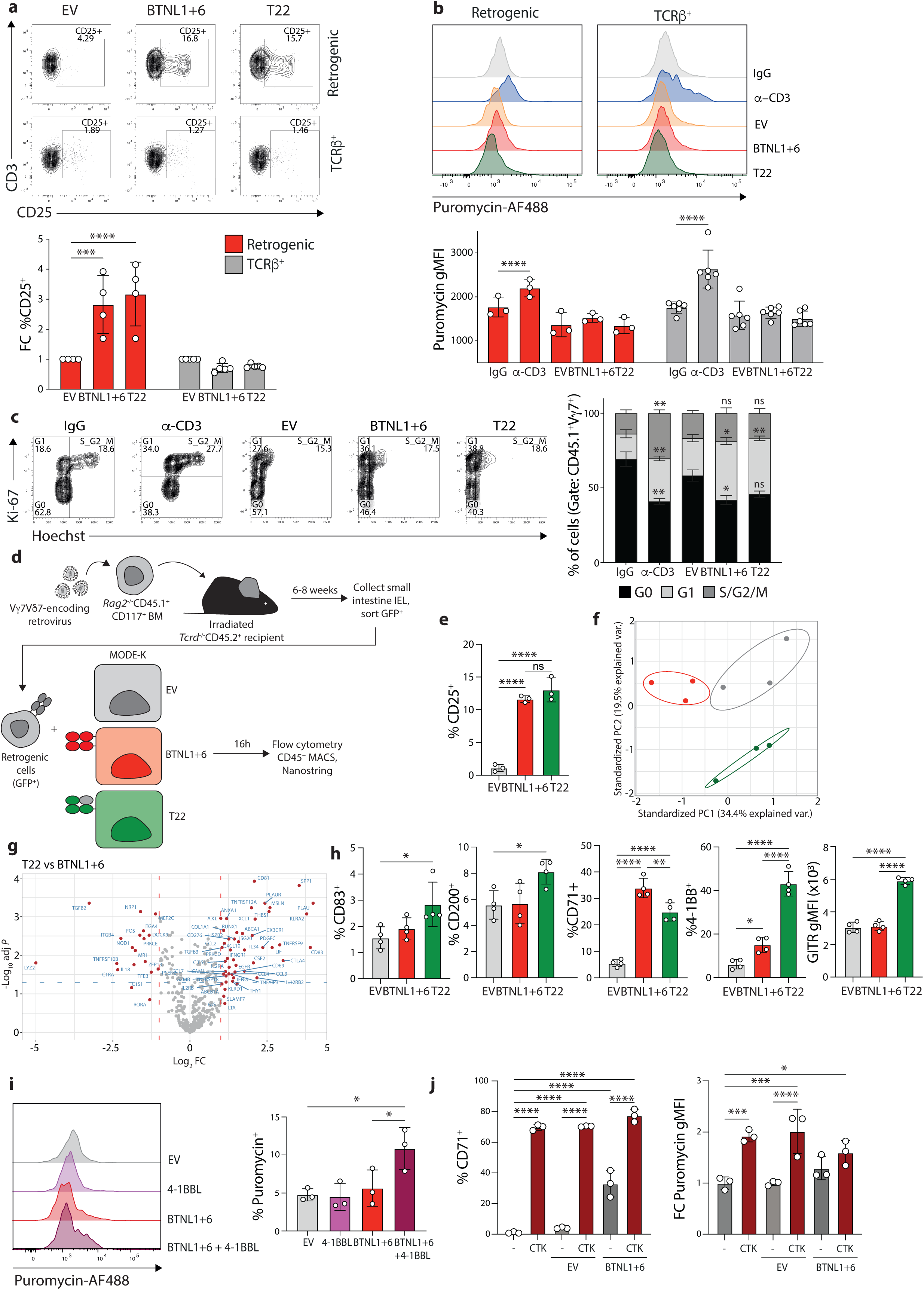
**a.** IEL were isolated from retrogenic mice and stimulated o.n. with MODE-K expressing an empty vector (EV), BTNL1+6 or T22. CD25 upregulation was analysed using flow cytometry. **b.** Puromycin incorporation was analysed by flow cytometry after o.n. stimulation of retrogenic IEL with plate-bound IgG or anti-CD3, or with MODE-K expressing EV, BTNL1+6 or T22. **c.** Analysis of cell cycle in activated Vγ7+ cells using Ki-67 and Hoechst staining. **d.** Experimental setup to analyse gene expression following stimulation of retrogenic cells. Retrogenic mice were generated as described, and small intestinal IEL isolated. CD45.1 and GFP positive cells were sorted via FACS and co-cultured for 16h with EV, BTNL1+6 or T22-expressing MODE-K cells. **e.** CD25 expression was analysed after stimulation using flow cytometry. **f.** The remainder of the cells was isolated using CD45 MACS beads and gene expression assessed using Nanostring. PCA analysis of overall gene expression. **g.** Volcano plot of genes differentially expressed between BTNL1+6 and T22-stimulated cells. **h.** IEL from retrogenic mice were isolated, stimulated o.n. with EV, BTNL1+6 or T22 MODE-K cells and assessed for CD83, CD200, CD71, 4-1BB and GITR expression via flow cytometry. **i.** WT IEL were stimulated o.n. with MODE-K expressing EV, 4-1BBL, BTNL1+6 or BTNL1+6+4-1BBL. Puromycin incorporation was analysed by flow cytometry. **j.** WT IEL were stimulated o.n. with IL-12, IL-18 and TNF-α (CTK), alone or in combination with MODE-K expressing EV or BTNL1+6. CD71 upregulation and puromycin incorporation were analysed by flow cytometry.

Nonetheless, to determine whether the two distinct modes of TCR engagement might elicit distinct functional programmes, we assessed global transcriptional responses. Retrogenic γδ T cells were sorted based on GFP expression and subsequently co-cultured with MODE-K cells expressing BTNL1+6, T22 or empty vector (Figure 4d). Following overnight stimulation, IELs were re-isolated and subjected to targeted transcriptional profiling using NanoString. In parallel, analysis of CD25 upregulation revealed highly comparable levels of activation in response to BTNL1+6 and T22 stimulation, respectively (Figure 4e).

Principal component analysis revealed clear segregation of EV, BTNL1+6, and T22-stimulated retrogenic γδ T cells: BTNL1+6-stimulated cells differed from T22-stimulated and EV cells primarily in PCA1, accounting for ∼35% of variance, whereas T22-stimulated cells differed from BTNL1+6-stimulated cells and EV cells primarily in PCA2 accounting for ∼20% of the variance (Figure 4f). Thus, taken together, our data demonstrated that primary IEL made largely comparable responses to BTNL1+6 and T22, but that nonetheless the transcriptional programmes induced by these two TCR ligands were distinct.

### Signal 2 and Signal 3 overcome attenuated TCR responses

Based on qualitative differences in the transcriptional responses elicited by the respective ligands and on the use of flow cytometry to assess changes in surface protein expression, T22 stimulation appeared to preferentially upregulate the fraction of cells expressing and/or the intensity of expression of genes and proteins associated with T cell co-stimulation and regulation, specifically CD83, CD200, TNFRSF18 (a.k.a. GITR) and TNFRSF9 (a.k.a. 4-1BB) (Figure 4g, h). In most cases, the percentages of cells affected were very low, and hence of uncertain biological significance. By contrast, T22 increased the fraction of 4-1BB^+^ cells relative to EV from ∼7% to >40%, while BTNL1+6 induced up to ∼20% of cells to be positive (Figure 4h; see also Figure 1a). Note that the comparable levels of CD25 upregulation, shown above (Figure 4e) and the increase in the expression of several genes and proteins, including CD71 in BTNL1+6-stimulated cultures relative to T22-stimulated cultures (Figure 4g, h) excludes that BTNL1+6-induced responses were simply of lower magnitude.

We hypothesised that the upregulation of 4-1BB might make IEL responsive to 4-1BB-ligand (4-1BBL) and that this could synergise with TCR signalling to achieve a more complete activation of responding IEL (Signal 2). To test this, we generated MODE-K cell lines expressing BTNL1+6 alone or in combination with 4-1BBL and, reverting back to IEL from WT C57BL/6 rather than retrogenic mice, we found that 4-1BBL facilitated, for the first time, a global increase in protein translation in BTNL1+6-stimulated primary IEL (Figure 4i). Thus, given that 4-1BBL would only rarely be expressed in healthy tissues that express BTNL1+6, it would be the absence of such co-stimulation that could limit IEL responses to TCR-mediated engagement. Conversely, 4-1BBL is often strongly expressed in inflammatory settings (28–31), where it could permit a more complete IEL response to stress-regulated adaptive ligands, such as T10/T22 (32).

In this same vein, it is well established that primary IEL can make effector responses, e.g., IFNγ release, to cytokines (Signal 3) such as IL-12, IL-18 and TNF-α (33,34), that again would be expressed in stressed and/or inflammatory settings. When primary IEL were co-cultured with either EV or BTNL1+6-expressing MODE-K cells in the presence of IL-12+IL- 18+TNF-α (CTK), the BTNL-induced increase in CD71 expression (see above) was increased still further, and global protein translation was increased by CTK alone and CTK plus BTNL1+6, but again not by BTNL1+6 alone (Figure 4j). In sum, the complete responsiveness of IEL is triggered by cytokines and co-stimulatory molecules together with TCR ligand engagement, but not by TCR ligand engagement alone.

### The requirements for TCRγδ-mediated responses to BTNLs

We next investigated the molecular requirements for the attenuated activation of IEL following TCR engagement, initially by adding a series of signal transduction inhibitors to retrogenic IEL stimulated with either BTNL1+6 or T22. Consistent with the phospho-flow data shown above (Figure 2), inhibitors of LCK greatly reduced CD25 upregulation in response to BTNL1+6 stimulation (Figure 5a; Supp Fig 3a). The same was true for PP2, a general SRC-kinase inhibitor, and flavopiridol, a broad CDK and transcriptional inhibitor (Figure 5a; Supp Fig 3a). Activation was slightly affected by a CD45 inhibitor, was strongly suppressed by inhibiting PI3Kδ, but was not affected by rapamycin, which ordinarily inhibits *Il2ra* gene transcription in addition to reducing mTOR activity and downstream cell responses (Figure 5a). The different inhibitors had broadly the same impacts on responses to T22 engagement (Figure 5b).

**Figure 5.**
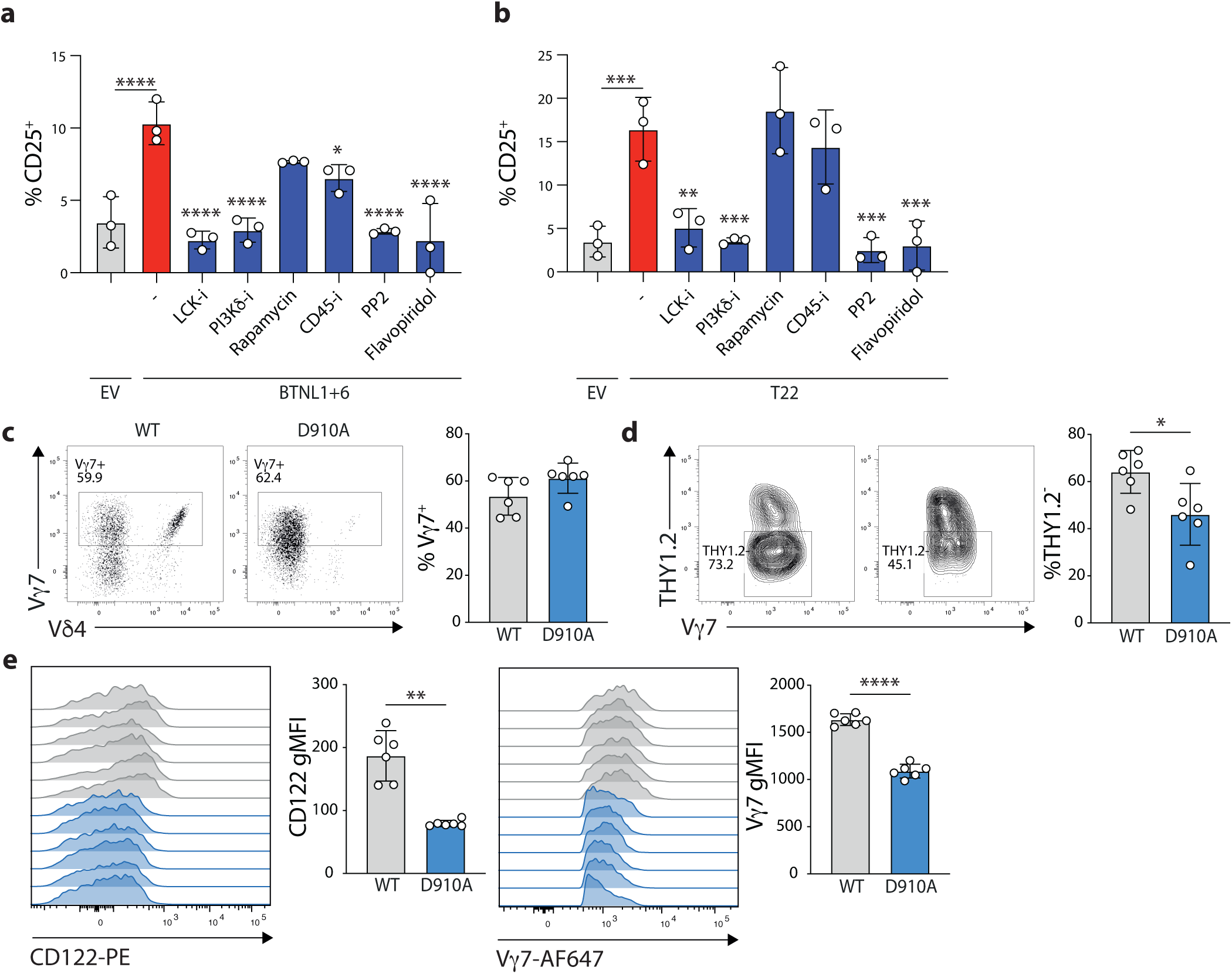
Retrogenic IEL were isolated and stimulated o.n. with MODE-K expressing EV, BTNL1+6 or T22, alone or in the presence of inhibitors targeting LCK, PI3Kd or CD45, rapamycin, PP2 or flavopiridol. (**a, b**) Quantification of the percentage of CD25+ cells following stimulation. IEL were isolated from WT or p110δ^D910A^ (D910A) mice and Vγ7+ cells (**c**) and maturation marker expression (**d**, **e**) assessed by flow cytometry.

Given that BTNL1+6 signalling was independent of rapamycin and failed to drive AKT/mTOR activation (see above, Figure 2), we further investigated components that might be related to this relative attenuation of TCR signal trasnsduction. Specifically, we investigated PI3Kδ that is a primary promoter of AKT/mTOR activation in leukocytes (35). First, we examined PI3Kδ D910A mice in which PI3Kδ is catalytically inactive (36). The IEL compartment of adult mice harboured Vγ7^+^ IEL, but there was an almost complete absence of Vγ7Vδ4^+^ IEL and a corresponding increase in Vδ6.3^+^ cells, both of which were previously observed in *Btnl6*-deficient mice(37) (Figure 5c; Supp Fig 3b). In examining other markers of BTNL-mediated Vγ7^+^ IEL selection, we noted that Vγ7^+^ IEL from D910A mice upregulated LAG3 as normal (Supp Fig 3c) but failed to downregulate THY1 to the normal degree and failed to upregulate CD122 and TCR expression to wild-type levels (Figure 5d, e). Finally, Vγ7^+^ IEL from D910A mice showed dysregulated CD25 induction relative to wildtype mice, although this was in part because of a higher background level of activation (Supp Fig 3d). These data, taken together with the impact of the inhibitor (above), establish PI3Kδ as crucially contributing to BTNL-induced signal transduction.

### Cell-intrinsic relief of attenuated TCRγδ-mediated responses

We next considered that the failure to activate AKT and mTOR might perhaps reflect weak PI3Kδ signalling downstream of BTNL-TCR interactions. To test this, we employed the PI3Kδ E1020K mouse in which PI3Kδ is constitutively active (38). These mice harboured typical numbers of Vγ7^+^ IEL, but unlike in D910A mice, IEL maturation was largely normal and possibly slightly exaggerated, at least as measured by CD122 upregulation (Figure 6a-c). Additionally, Vδ usage was comparable to wild-type mice (Figure 6d). Nonetheless, enforced PI3Kδ activity conspicuously failed to restore either puromycin incorporation or increase activation marker expression following BTNL1+6 stimulation relative to wild type mice (Figure 6e,f).

**Figure 6.**
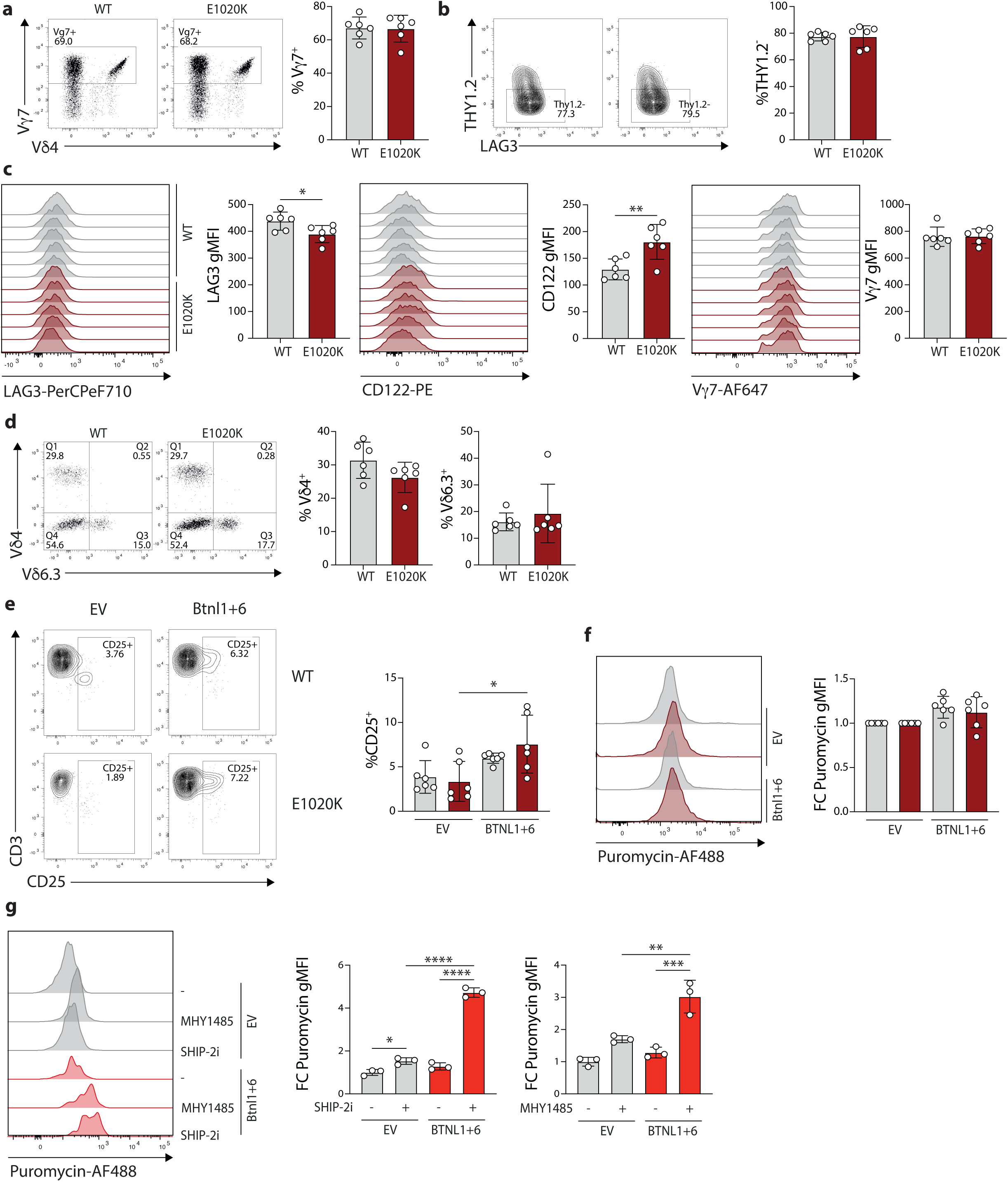
IEL were isolated from WT or *Pik3cd*^E1020K/+^ (E1020K) mice and Vγ7+ cells (**a**), maturation marker expression (**b**, **c**) and Vdelta chain usage (**d**) assessed by flow cytometry. IEL were stimulated o.n. with MODE-K expressing EV or BTNL1+6 and (**e**) CD25 upregulation and (**f**) puromycin incorporation were analysed by flow cytometry. **g.** WT IEL were stimulated o.n. with MODE-K expressing EV or BTNL1+6, alone or in the presence of MHY1485 or SHIP-2 inhibitor (SHIP-2i). Puromycin incorporation was measured by flow cytometry.

These findings suggested the presence of an additional regulatory checkpoint that specifically neutralised the impact of enforced PI3Kδ activity. Stimulating IEL in the presence of the SHP-2 protein phosphatase inhibitor SHP099 did not reveal a significant increase in translation rate following BTNL1/6 stimulation (Supp Fig 4a). We next focused on lipid phosphatases, which act proximally on PI3K-generated second messengers (35,39). Indeed, SHIP-1 (a.k.a Inpp5d) is expressed at noticeably high levels by γδ IEL and, albeit to a lesser extent, SHIP-2 (a.k.a. Inpp1) is also highly expressed (15,40), particularly in TCRγδ^+^ IEL and in CD8αβTCRαβ^+^ IEL relative to CD8αα^+^TCRαβ^+^ IEL and lymph node-derived naïve CD8 T cells (5). Inhibition of SHIP-1 had no effect on BTNL1/6-induced translation (Supp Fig 4b), whereas inhibition of SHIP-2 significantly increased puromycin incorporation following BTNL1/6 stimulation (Figure 6g). Moreover, puromycin incorporation was also significantly increased by MHY1485, an mTOR agonist (Figure 6g). These results identify SHIP-2 as a selective negative regulator constraining cellular reprogramming downstream of BTNL1/6 engagement, most likely through its suppression of mTOR activation.

## Discussion

Body surfaces must be able to maintain semi-permeable barriers that simultaneously allow exchange with the outside world, including nutrient uptake and water exchange, while limiting the entry of pathogens. Part of this is orchestrated by IEL, which appear replete with cytotoxic effectors, chemokines and epithelial growth regulators. Clearly, it is crucial for IEL to respond effectively when required, while minimising damage under homeostatic conditions. Indeed, dysregulation of the IEL compartment is a feature of several autoimmune or autoinflammatory conditions, including IBD and celiac disease (34,41,42).

TCRγδ is key to the regulation of γδ IEL (14), in which regard a striking finding over the past decade has been that the TCRs of human and mouse gut γδ IEL can deploy two modes of ligand engagement: a largely innate recognition that employs TCRVγ motifs to bind to BTNL molecules expressed by healthy epithelium (6)(17)(19), and an adaptive, CDR3-dependent recognition of “stress-associated” molecules that are commonly MHC related, e.g., human CD1c and EPCR, and mouse T10/T22 (18,24,43). An IEL’s interpretation of TCR engagement of these distinct ligand classes must be appropriate to the setting, specifically, a constrained response within healthy tissue and a robust response to tissue dysregulation, but how this distinction occurs has remained unelucidated.

To investigate this, we have studied primary murine IEL, consistently observing that Vγ7^+^ IEL (the majority of γδ IEL) are competent to respond conventionally to anti-CD3, but that the same cells show atypical, selectively attenuated responses to BTNL1+6 engagement. First, they fail to activate ZAP70, a trait which was recently shown to attenuate TCR signalling in neonatal thymocytes, allowing them to avoid negative selection. This may explain the hitherto unresolved observation that gut IEL are a haven for autoreactive T cells (10).

Attenuated ZAP70 and SYK activation downstream of the TCR may explain the failure of BTNL engagement to promote even transient AKT and mTOR activation, thereby limiting IEL proliferation and protein translation. Initially, we were surprised to find that Vγ7^+^ IEL likewise made attenuated responses to a clonotypic ligand (T22) that, unlike BTNLs, engages the TCR in a CDR3-dependent manner and is associated with cell and tissue dysregulation. Indeed, among the few differences that we could identify in the IEL responses to the two classes of ligand was the greater upregulation of 4-1BB by the clonotypic ligand. Future studies should expand on the mechanistic basis for this difference, which may include conformational changes to the TCR in response to ligation of the BTNL-binding face or the CDR3s (44), respectively, and/or differential receptor clustering and exclusion or inclusion of specific co-stimulatory receptors at the immunological synapse.

Irrespective of mechanism, TCR signalling will permit IEL to respond strongly to 4-1BBL engagement, which was apparent in the upregulation of global protein synthesis induced by TCR stimulation in the presence of 4-1BBL. Since 4-1BBL is negligibly expressed in the healthy gut, such transition to full-blown IEL activation would not occur, whereas it would be relevant for IEL engaging clonotypic ligands associated with tissue dysregulation. Hence, a feed-forward loop initiated by the TCR (4-1BB upregulation) and compounded by 4-1BBL expression would permit appropriately distinct IEL responses to healthy and dysregulated tissue, respectively. The commitment to full-blown activation is also facilitated by inflammatory cytokines that would also be robustly expressed in dysregulated but not healthy epithelia.

Our studies have also attributed a function to SHIP-2 which is differentially expressed by γδ IEL and lymph node-derived T cells (40). Specifically, its inhibition derepressed the TCR-mediated response to BTNL1+6. This would be consistent with its preventing accumulation of PI(3,4,5)P3 induced by PI3K, thereby constraining AKT and mTOR activation (35,39). Aligned with this, enforced mTOR activation also derepressed the response to BTNL1+6. Presumably, the suppression of AKT and mTOR activation downstream of TCR engagement can be overcome by co-stimulation either by virtue of enforced PI3K activation, that out-titrates SHIP-2, and/or by indirect mTOR activation via TRAF signalling, as is the *modus operandi* of 4-1BB and other TNFRSF co-stimulators (22). Inflammatory cytokines may likewise induce proliferation and global protein synthesis *via* STAT activation, but again those effects are γδ TCR-dependent (14).

In sum, the γδ TCR signals atypically in primary IEL responding to both innate and adaptive ligands. Our findings show that a cell-intrinsic contribution of this is mediated by SHIP-2, thereby creating a foundation on which cell-extrinsic co-stimulators and cytokines can act to tune IEL responses to TCR engagement according to context. Whether SHIP-2 activity itself is tunable in γδ T cells remains unknown. In other cellular settings, immunoreceptors such as FcγRIIB can recruit and position SHIP-2 within receptor-associated signalosomes (45,46). SHIP-2 activity and phosphorylation is also responsive to growth factors and insulin signalling (47,48). These findings raise the possibility that SHIP-2 could integrate environmental signals to selectively modulate γδ T cell activity. At the same time, we have been able to offer no functional explanation for the considerable enrichment of SHIP-1 in IEL. Moreover, our findings are unlikely to have described the complete picture, given that there are several atypical aspects to IEL signal transduction, as considered in this paper’s introduction. Certainly, many further traits of IEL remain to be resolved, including their non-canonical CD3 composition, how signals from activating and inhibitory receptors are integrated by the cells and the organising principles governing ligand recognition. A better understanding of these may permit an informed approach to IEL dysregulation in conditions such as celiac disease and IBD.

## Methods

### Mice

Wild-type C57BL/6J, *Nur77*-eGFP, *Btnl1*^-/-^ and *B2m*^-/-^ mice were maintained at The Francis Crick Institute’s animal facility under SPF conditions and used for experiments between 6 and 12 weeks of age. Animals were maintained under a 12h light/dark cycle and access to food and water *ad libitum*. PI3Kδ^D910A^ (36) and PI3Kδ^E1020K^ (38) mice were bred and maintained at the University of Cambridge animal facility and used between 5 and 11 weeks of age. All animal work was carried out in compliance with the UK Home Office regulations under project licenses PP5000699 to ACH and PPL P0AB4361E.

### IEL isolation

Mouse IELs were collected from the small intestine. Briefly, the small intestine was excised, cut longitudinally, washed twice with PBS and cut into 0.5cm pieces. These were placed in 10mL of complete RPMI supplemented with 1mM DTT and incubated at RT for 20min in a rotating incubator. Tissue was then centrifuged and resuspended in fresh complete RPMI, vortexed twice and passed through 70um cell strainers. IEL were isolated by centrifugation in an 80:40:20 Percoll gradient and cultured or processed for flow cytometry.

### IEL stimulation

For IEL stimulations, 1×10^5^ MODE-K cells expressing an empty vector (EV), BTNL1+6, 4-1BBL or 4-1BBL in combination with BTNL1+6 were plated onto 48-well plates 24h prior to IEL isolation. The following day, 1.5×10^5^ IEL resuspended in 200uL of RPMI supplemented with L-glutamine, 10% HI FBS, 1% Pen/Strep, 10mM HEPES, 1mM Sodium Pyruvate, 1x NEAA, 50uM β-mercaptoethanol, IL-2 10U/mL, IL-15 10ng/mL, IL-3 100U/mL and IL-4 200U/mL were overlaid onto the monolayer. Cells were typically co-cultured overnight (16-18h) at 37C with 5% CO2. For phospho-flow analysis, IELs were rested in complete RPMI containing 2.5% FBS and incubated for the indicated times with MODE-K cells, or IgG/anti-CD3 in complete RPMI. Following stimulation, cells were fixed immediately for 10min at RT with 1.5% PFA, before washing and fixation with ice-cold 100% methanol.

For antibody stimulations, 5ug/mL of IgG or anti-CD3 (2C11) were plated in PBS on 48-well plates for 4h prior to IEL isolation. Plates were incubated at 37C until use and washed twice with PBS before addition of the IEL suspension.

For cytokine stimulations (CTK), 100ng/mL of IL-12, 100ng/mL IL-18 and 50ng/mL TNF-α were added to the culture.

### Puromycin incorporation assay

Following overnight stimulations, IEL were pulsed with 10ug/mL puromycin for 10 minutes at 37C. Cells were then harvested and analysed by flow cytometry.

### Flow cytometry

Cell suspensions were stained for 15min at RT with Live/Dead dyes (Invitrogen), followed by 30min of staining at 4C with membrane antibodies (see Table 1) and Fc block. For intracellular staining, cell suspensions were fixed and permeabilised with BD Cytofix/Cytoperm, followed by BD Perm/Wash according to the manufacturer’s instructions. Cells were then incubated with anti-Puromycin or anti-Ki-67 for 60 min at RT before analysis. For analysis of cell cycle, cells were stained with Hoechst for 10min immediately before acquisition. Alternatively, for phosphoflow, following methanol fixation, cells were washed twice and stained with appropriate combinations of directly conjugated primary antibodies, or primary and secondary antibodies (see Table 1).

**Table 1.** Antibodies used for flow cytometry.

| Antibody | Clone | Source | Cat no. | Dilution |
| --- | --- | --- | --- | --- |
| CD3-APCCy7 | 17A2 | BioLegend | 100222 | 1:100 |
| CD122-PE | TM-β1 | BioLegend | 123209 | 1:100 |
| THY1.2-BV510 | 53-2.1 | BioLegend | 140319 | 1:100 |
| LAG3-PerCPeF710 | C9B7W | eBioscience | 46-2231-80 | 1:100 |
| TCR Vd4-PE | GL2 | BioLegend | 134905 | 1:100 |
| TCR Vg7 | F2.67 | Institut Pasteur, Paris, P. Pereira | N/A | 1:400 |
| CD25-PerCPCy5.5 | PC61 | BioLegend | 102030 | 1:100 |
| CD71-PECy7 | RI7217 | BioLegend | 113812 | 1:100 |
| 4-1BB (CD137)-PE | TKS-1 | BioLegend | 107105 | 1:200 |
| Puromycin-AF488 | 12D10 | Sigma | MABE343-AF488 | 1:100 |
| Puromycin-AF647 | 12D10 | Sigma | MABE343-AF647 | 1:100 |
| Ki-67-FITC | SolA15 | Invitrogen | 11-5698-82 | 1:100 |
| pCD3z (Y142)-PE | K25-407.69 | Fisher Scientific | 15819398 | 1:50 |
| p-LCK (Y505) |  | Santa Cruz Biotechnology | sc-136184 | 1:50 |
| p-LCK (Y394) | Rabbit Polyclonal | Antibodies Online | ABIN3182460 | 1:50 |
| p-ZAP70 (Y493) | N3kobu5 | eBioscience | 46-9006-41 | 1:50 |
| p-mTOR (S2448)-PECy7 | MRRBY | eBioscience | 25-9718-41 | 1:50 |
| p-SYK (Y348)-PE | Moch1ct | Invitrogen | 12-9014-42 | 1:50 |
| p-AKT (S473) | 193H12 | Cell Signalling | 4058 | 1:50 |
| p-ERK1/2 | D13.14.4E | Cell Signalling | 4370S | 1:50 |
| Goat anti-Rabbit IgG-AF488 |  | Invitrogen | A-11008 | 1:50 |
| IkBα-PE | MFRDTRK | eBioscience | 12-9036-42 | 1:50 |
| CD45.1 | A20 | BioLegend | 110741 | 1:100 |
| CD83-PE | Michel-19 | BioLegend | 121507 | 1:200 |
| CD200-PE | OX-110 | BioLegend | 123907 | 1:200 |
| GITR (CD357)-APC | DTA-1 | BioLegend | 126311 | 1:100 |
| Vd6.3-BV711 | 8F4H7B7 | BD | 744476 | 1:100 |
| CD45-AF700 | 30-F11 | BioLegend | 103128 | 1:100 |
| TCRβ-BV605 | H57-597 | BD | 562840 | 1:100 |
| TCRβ-BV510 | H57-597 | BD | 563221 | 1:100 |

Samples were acquired in an LSR Fortessa and analysis was performed using FlowJo (Version 10.10.0).

### Retrogenics

Retrovirus was generated by transfection of Plat-E packaging cells with pMIG-Vγ7Vδ7 vector encoding a BTNL1+6 and T22 dual-reactive TCR. Viral supernatants were collected 48h post-transfection, filtered and concentrated by centrifugation at 20,000xg for 2h. Legs were harvested from *Rag2*^-/-^CD45.1+ donors and HSCs were enriched via MACS using CD117 beads (Miltenyi), according to the manufacturer’s instructions. Cells were transferred to Stempan medium (Stem Cell Technologies) supplemented with SCF, hIL-6, TPO, Flt3L and IL-3 and spinfected for 1.5h at 2300rpm in a retronectin-coated plate.

Recipient mice were irradiated with a split-dose of 11Gy. The following day, 1×10^6^ transduced CD117+ cells were transplanted i.v. into the mice, and allowed to reconstitute for 6-8 weeks.

### Nanostring transcriptome analysis

Live GFP^+^ cells were FACS sorted from retrogenic mice IEL and co-cultured overnight in triplicate with MODE-K expressing EV, BTNL1+6 or T22. Following the incubation, retrogenic cells were re-isolated using MACS CD45 beads (Miltenyi) and lysates prepared from 30,000-50,000 cells per sample. Lysates were hybridised to the nCounter PanCancer Mouse Immune Profiling Panel (Nanostring Technologies) to evaluate expression of 770 targets. Hybridised samples were processed on an nCounter prep station and data were collected on an nCounter digital analyser (Nanostring), according to the manufacturer’s instructions. Raw data were imported into nSolver4.0 (Nanostring) for quality checks, thresholding and normalisation. Differential expression analysis was carried out using R-4.1.1, using functions from the limma package. Statistical significance was calculated using a moderated t-test with Benjamini-Hochberg correction for multiple comparisons (false discovery rate). All plots were produced using variations of the ggplot2 package.

### Inhibitor treatment

Dependence on specific kinases and phosphatases for signalling was studied by carrying out overnight stimulation studies in the presence of Lck inhibitor (A-770041) 1μM, PI3Kd inhibitor (IC-87114) 10μM, Rapamycin 50nM, CD45 inhibitor (SC-222223, Insight Biotechnology) 10μM, Src inhibitor (PP2) 10μM, Flavopiridol 1μM, SHP-2 inhibitor (SHP099) 2.5μM, mTOR agonist (MHY1485) 10μM, SHIP-2 inhibitor (AS1938909) 10μM or SHIP-1 inhibitor (3AC) 7.5μM.

### Immunohistochemistry

After fixation for 24 hours in 10% neutral buffered formalin, mouse small intestines were processed on a Tissue-Tek VIP 6 AI processor for embedding in paraffin wax. Tissue sections were cut at 3μm for downstream staining.

For immunohistochemistry, a Discovery Ultra IHC slide staining system (Roche) was used. Tissue sections were incubated with anti-GFP (1:250, Abcam, ab6673) followed by incubation with the Omnimap anti-Goat HRP (RUO) (Roche). Immunodetection was performed using DABMap Kit (Roche), and finally sections were counterstained in Haematoxylin.

## Acknowledgments

We thank the Francis Crick Institute BRF facility, and in particular C. Watkins, L. Valencia and P. Owers for technical support; the Flow Cytometry facility, with special thanks to A. Agua-Doce for assistance with cell sorting; the Experimental Histopathology Facility and the Genomics Facility. A.C.H. received support from the Francis Crick Institute (FC001093), which receives core funding from Cancer Research UK, the Medical Research Council (MRC) and the Wellcome Trust. L.M. received support from the Academy of Medical Sciences Springboard Award (SBF009/1064).

## Ethics Declarations

Adrian Hayday is a consultant for Ferring Ventures, TransImmune AG, Prokarium, Avidicure, and IMUBiosciences and has equity holdings in TransImmune AG, IMUBiosciences, Prokarium, and HS Lifesciences. None of these entities contributed in any way to the conceptualisation or content of this study.

## Figure legends

**Supplementary Figure 1**

WT IEL were co-cultured o.n. with plate-bound IgG or anti-CD3, or with MODE-K cells transduced with empty vector (EV) or BTNL1+6, in samples matched to Figure 1c and d. IEL activation was measured by analysing CD25 on Vγ7+ cells.

**Supplementary Figure 2**

Tissue localisation of retrogenic Vγ7 T cells was analysed by GFP immunohistochemistry of FFPE small intestine sections. Arrows highlight examples of GFP+ cells. Scale bar: 100μm

**Supplementary Figure 3**

Retrogenic IEL were isolated and stimulated o.n. with MODE-K expressing EV, BTNL1+6 or T22, alone or in the presence of inhibitors targeting LCK, PI3Kd or CD45, rapamycin, PP2 or flavopiridol. Representative flow cytometry plots of CD25 upregulation are shown in (**a**). IEL were isolated from WT or p110δ^D910A^ (D910A) mice and Vdelta chain usage (**b**) and LAG3 expression (**c**) were assessed by flow cytometry. **d.** IEL were stimulated o.n. with MODE-K cells expressing EV or BTNL1+6 and CD25 upregulation was analysed by flow cytometry.

**Supplementary Figure 4**

WT IEL were stimulated o.n. with MODE-K expressing EV or BTNL1+6, alone or in the presence of SHP099 (SHP-2 inhibitor) (**a**) or SHIP-1 inhibitor (SHIP-1i) (**b**). Puromycin incorporation was measured by flow cytometry.

