## Supplementary figures and images for "Atypical T cell receptor signalling permits distinct gut γδ T cell responses to healthy versus dysregulated tissues"

### Supplemental Figures

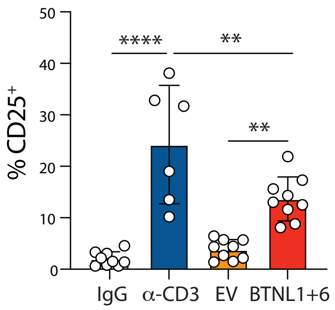


**Supp Figure 1**

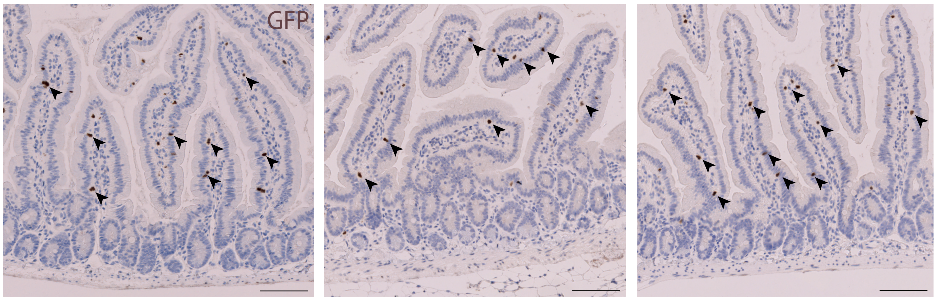


**Supp Figure 2**

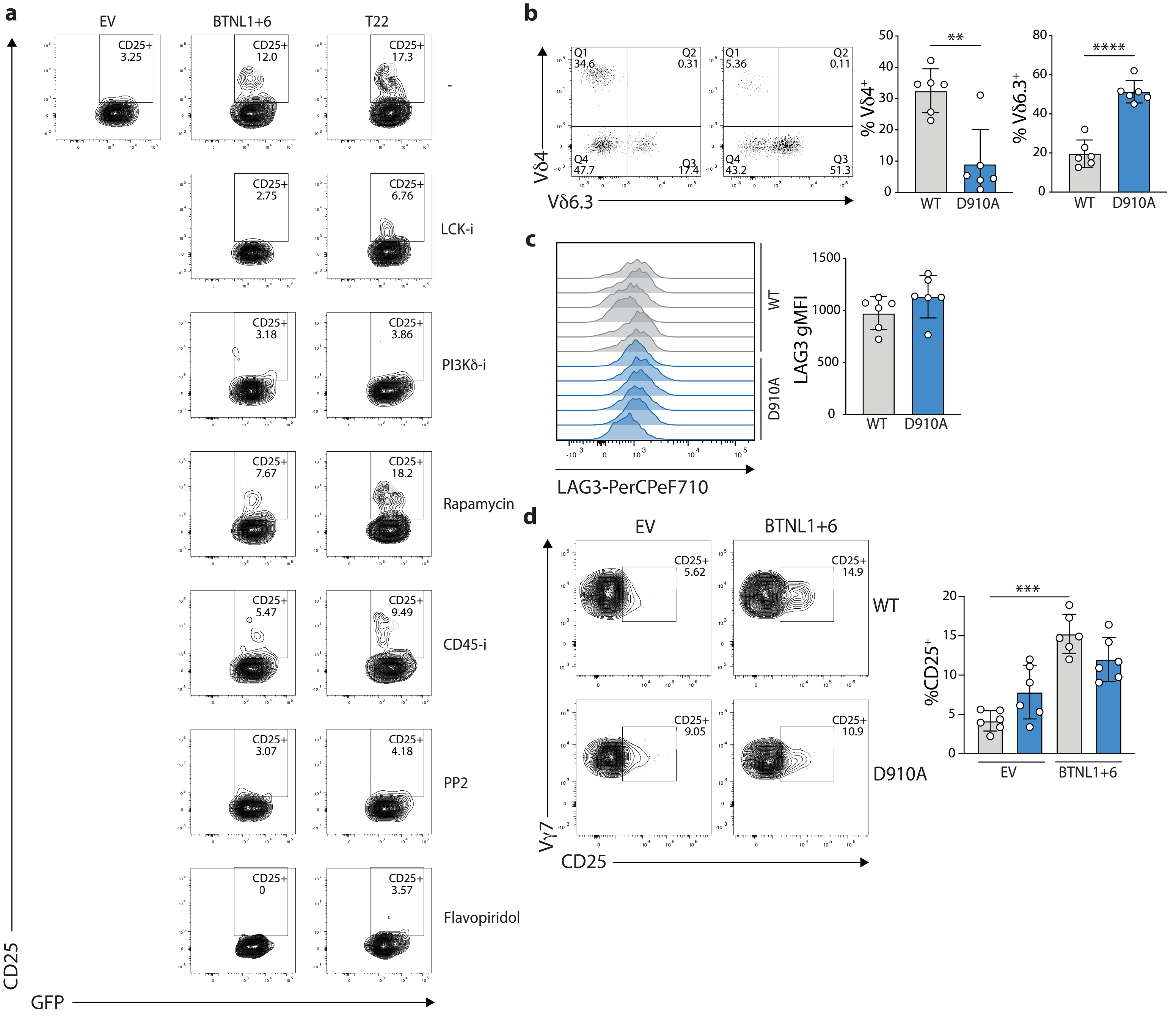


**Supp Figure 3**

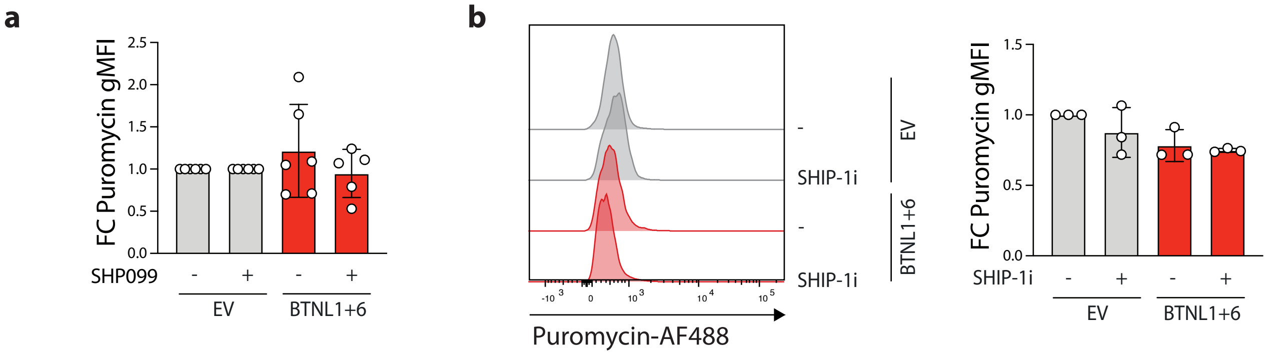


**Supp Figure 4**
